# The autophagy protein Atg9 functions in glia and contributes to parkinsonian symptoms in a *Drosophila* model of Parkinson’s disease

**DOI:** 10.1101/2023.04.03.535355

**Authors:** Shuanglong Yi, Linfang Wang, Margaret S. Ho, Shiping Zhang

**Affiliations:** School of Life Science and Technology, ShanghaiTech University, Shanghai, 201210, China; University of Chinese Academy of Sciences, Beijing, 100049, China

**Author notes:** Corresponding author: Shiping Zhang, Ph.D.

**Keywords:** Glia, Atg9, Parkinson’s disease, Autophagy

## Abstract

Parkinson’s disease (PD) is a progressive neurodegenerative disorder characterized by the motor deficits, selective loss of dopaminergic (DA) neurons, and the brain accumulation of α-synuclein (α-syn)-composed protein aggregates called Lewy bodies (LBs). Whereas dysfunction in the protein degradation pathway, like autophagy in neurons, has been demonstrated as a critical mechanism for eliminating protein aggregates in PD, how protein aggregates are eliminated in the other brain cell type, glia, is less well characterized. In the present study, we show that Atg9, the only transmembrane protein in the core autophagy pathway, is highly expressed in *Drosophila* adult brain glia. Results from immunostaining and live-cell imaging analysis reveal that a significant portion of Atg9 localizes to the trans-Golgi network (TGN), autophagosomes, and lysosomes in glia; Atg9 is persistently in contact with these organelles. Lacking glial *atg9* reduces the number of omegasome and autophagosome and impairs autophagic substrate degradation, suggesting that glial Atg9 participates in the early steps of autophagy, hence the control of autophagic degradation. Importantly, loss of glial *atg9* induces parkinsonian symptoms in *Drosophila* including progressive DA neuron loss and locomotion deficits. Our findings identify a functional role of Atg9 in glial autophagy and establish a potential link between glial autophagy and PD. These results provide new insights on the underlying mechanism of PD.

## Introduction

Parkinson’s disease (PD), first described by James Parkinson in 1817 [1], is the second most common neurodegenerative disorder mainly affecting elderly people [2]. Clinically, PD patients display both motor symptoms (e.g. rest tremor, rigidity, bradykinesia, and postural instability) and non-motor symptoms (e.g. depression, sleep disorders, anosmia, and cognitive impairment) [3]. The patient brains exhibit progressive loss of dopaminergic (DA) neurons in the *substantia nigra pars compacta* (*SNpc*) and the accumulation of Lewy bodies (LBs) and Lewy neurites (LNs) composed of α-synuclein (α-syn) [3-5]. Whereas accumulating evidence has suggested that neurons are the central players for these neuropathological hallmarks in PD, glia, more than mere bystanders that simply support neurons, are also activated in the patient brains and contribute significantly to PD progression [6-9], thus increasing attention have been drawn to unravel the regulatory mechanisms of glia in PD.

Autophagy, a conserved self-eating process involving the engulfment of cytoplasmic materials or organelles, has been shown to protect the brain from neurodegeneration by eliminating toxic protein aggregates and defective mitochondria; impaired autophagy contributes to the underlying mechanism of neurodegenerative diseases such as Alzheimer’s disease (AD), PD, and Amyotrophic lateral sclerosis (ALS) [10-12]. Autophagy begins with the formation of phagophore from an isolated membrane, then the sealing of phagophore into a double-membrane autophagosome. Autophagosome subsequently fuses with lysosome for the degradation of the engulfed contents by lysosomal hydrolases in an acidic milieu. The resulting breakdown products are released and recycled back to the cytosol. The autophagy process is regulated by complexes composed of different autophagy-related (ATG) proteins including the Unc-51-like kinase 1 (ULK1) complex, the class III PI3K (PI3KC3) complex, the PI3P effectors, the Ubiquitin (Ub)-like conjugation system ATG12∼ATG5-ATG16L, and the ATG9 membrane delivery protein [13-16]. Autophagy occurs in basal conditions to maintain homeostatic function, and is upregulated upon starvation or cellular stress to supply amino acids or energy [13-16]. ATG9, the only known transmembrane protein in the core autophagy machinery, was first identified in yeast [17, 18]. ATG9 is a multi-spanning membrane protein and is proposed to be the membrane source of autophagosomes [19, 20]. ATG9 has divergent localizations in yeast and mammals. In yeast, Atg9 is localized on the cytoplasmic mobile vesicles (Atg9 vesicles) derived from the Golgi apparatus. During starvation, several Atg9 vesicles assembles into the pre-autophagosomal structure (PAS, also known as the phagophore-assembly site), and eventually incorporates onto the outer membrane of autophagosomes. After autophagosome formation, Atg9 is recycled back to the cytoplasm [21, 22]. Thus, Atg9 undergoes dynamic movements between the peripheral vesicular pools and PAS/autophagosomes. In mammals, ATG9 cycles between the trans-Golgi network (TGN), late endosomes, and plasma membrane under normal conditions. Upon amino acid starvation or autophagy induction, ATG9 disperses into peripheral compartments, endosomal membranes, and autophagosomes [23-25]. During autophagy, ATG9 traffics through multiple organelles, including recycling endosomes, to promote autophagy initiation and progression [24, 26]. However, ATG9 interacts dynamically and transiently with phagophores and autophagosomes without being incorporated onto them [24].

Interestingly, ATG9 dysregulation has been implicated in PD. α-syn overexpression causes ATG9 mislocalization and decreases omegasome formation by inhibiting Rab1a [27]. Recently, ATG9 is reported to transport in vesicles and undergo activity-dependent exo-endocytosis at the presynaptic sites in *C. elegans* [28]. Mutation in *unc-26/synaptojanin 1*, a gene associated with early-onset Parkinsonism, disrupts endocytosis and results in abnormal ATG9 accumulation at clathrin-rich synaptic foci, leading to defects in activity-induced presynaptic autophagy [28]. Nevertheless, the exact role of ATG9 in PD and whether it acts differently in neurons or glia has not been investigated yet. Here we present evidence that the *Drosophila atg9* is highly expressed in glial cells. By immunostaining and live-cell imaging analysis, Atg9 is detected on TGN, autophagosomes, and lysosomes in glia, and persistently in contact with these organelles. Knocking down *atg9* expression in glia by RNA interference (RNAi) reduces the number of omegasome and autophagosome and impairs autophagic substrate degradation, suggesting that glial Atg9, like Atg9 in other tissues, regulates early steps of autophagy. Importantly, lack of glial *atg9* induces parkinsonian symptoms including progressive DA neuron loss and locomotion defect. Thus, our findings uncover the autophagic role of Atg9 in glia contributing to PD; these results will provide new insights on developing therapeutic strategies for PD.

## Materials and Methods

### *Drosophila* genetics

Flies were raised at 25L with 70% humidity, and all crosses were carried out at 25°C. Fly strains acquired from Bloomington *Drosophila* Stock Center (BDSC) and Vienna *Drosophila* RNAi Center (VDRC) include *w*^*1118*^ (BL5905), *UAS-LacZ* (BL1777), *UASp-GFP*.*Golgi* (BL31422), *UASp-RFP*.*Golgi* (BL30908), *UASp-mCherry*.*Atg8a* (BL37750), *UAS-atg9-RNAi* (BL28055), and *UAS-atg9-RNAi*^#2^ (V10045). Strains described previously include: *repo-GAL4* [29], *10xUAS-IVS-Syn21-GFP-p10* [30], *UAS-Lamp1*.*GFP* [31], and *UAS-GFP*.*ZFYVE1* [32]. 3- or 10-day-old adult fly brains were dissected. Detailed fly genotypes are listed in Supplementary Materials.

### Molecular biology

The DNA sequences of Atg9 and Lamp1 were synthesized and subcloned into *pUAST-attB* vector containing a EGFP or a mCherry epitope tag by Genscript (Nanjing, China). Fly microinjection was conducted by the *Drosophila* Core Facility, Institute of Biochemistry and Cell Biology, Chinese Academy of Sciences.

### qRT-PCR

Total RNA was extracted from adult fly heads (about 20) using TransZol Up (Cat. #ET111-01, TransGen, Beijing, China) according to the manufacturer’s instructions. And HiScript III RT SuperMix (Cat. #R323-01, Vazyme, Nanjing, China) was used to reverse-transcribe cDNAs. qRT-PCR was carried out on ABI 7500 RT-PCR system using ChamQ Universal SYBR qPCR Master Mix (Cat. #Q711-02, Vazyme). ΔΔCT method was used for analysis, and *rp49* was chosen as control gene. Results are representative of at least three biological replicates.

Primers used are listed below:

*rp49*-F: CCACCAGTCGGATCGATATGC

*rp49*-R: CTCTTGAGAACGCAGGCGACC

*atg9-*F: AGCAGAAGCACGGATTCACA

*atg9-*R: GCAGTGCATCACAAAGGCAA

### FACS analysis

Adult fly brains were dissected in Schneider’s medium (Cat. #21720024, Invitrogen) and dissociated in PBS containing 0.18 units/ml fresh papain (Cat. #LK003178, Worthington, Freehold, NJ, USA) and 2.5mg/ml liberase TM (Cat. #5401119001, Roche, Basel, Switzerland). For qRT-PCR analysis, approximately 100 brains were dissected. Before use, papain was activated for 15 minutes at 37°C. After 1 hour of dissociation, cold Schneider’s medium was added to inactivate the enzyme. After filtering through a cell strainer (Cat. #352340, BD Falcon, Franklin Lakes, NJ, USA) into a flow cell tube (Cat. #352003, BD Falcon), samples were centrifuged and re-suspended in Schneider’s medium. The suspension was then kept on ice before being sent to the flow cytometer (FACSMelody, BD). To sort adult fly glia, fluorescent GFP signals were monitored.

### Immunohistochemistry

Adult fly brains were dissected and fixed in 4% formaldehyde for 35 minutes, washed three times in PBT (PBS + 0.1% TX-100), and then further dissected in PBS solution. Brains were then blocked in PBT solution containing 5% Normal Donkey Serum (NDS), stained with primary antibodies at 4°C overnight, and with secondary antibodies at room temperature for 2 hours. Primary antibodies used are: rabbit anti-Atg9 (1:200, Cat. #NB100-98679, Novus Biologicals, Littleton, CO, USA), mouse anti-NC82 (DSHB, 1:100, Cat. #NC82), rabbit anti-Ref(2)P (1:500, Cat. #ab178440, Abcam, Cambridge, UK), and rat anti-DDC (1:300, gift from Jay Hirsh). Secondary antibodies used were from Jackson ImmunoResearch (West Grove, PA, USA): donkey anti-rabbit Cy3 (1:1000, Cat. #711-166-152), donkey anti-rat Cy3 (1:1000, Cat. #712-165-153), and donkey anti-mouse Cy5 (1:500, Cat. #715-175-150).

### Western blot

Fly adult heads (50/genotype) were homogenized in lysis buffer (0.4% NP-40, 0.2 mM EDTA, 150 mM NaCl, 20% glycerol, 100 mM Tris-HCl pH7.5, 2% Tween 20, 0.5 mM phosphodiesterase inhibitors, and 1 mM PMSF) using a motorised pestle (Cat. #116005500, MP Biomedicals, Irvine, CA, USA). SDS-PAGE gels were used to separate proteins, which were then transferred to PVDF membranes (Cat. #IPFL00010, Millipore, Billerica, MA, USA). The membranes were blocked with 5% fat-free milk diluted in PBS with 0.1% Tween-20 for 40 minutes. Primary antibodies were incubated at 4°C overnight, followed by HRP-conjugated secondary antibodies at room temperature for 2 hours. Primary antibodies used are: rabbit anti-Atg9 (1:200, Cat. #NB100-98679, Novus Biologicals), rabbit anti-Ref(2)P (1:1000, Cat. #ab178440, Abcam), mouse anti-Mono- and poly-ubiquitinylated conjugates antibody (FK2, 1:500, Cat. #BML-PW8810, Enzo Life Sciences), and mouse anti-α-Tubulin (1:5000, Cat. #T9026, Sigma). Secondary antibodies used are from Jackson ImmunoResearch: goat anti-mouse-HRP (1:5000, Cat. #115-035-003), goat anti-rabbit-HRP (1:5000, Cat. #111-036-003). Bands were visualized by Clarity Western ECL Substrate (Cat. #WBKLS0500, Millipore). Results are representative of at least three biological replicates.

### *In-vivo* live-cell imaging analysis

Live adult fly brains were dissected and kept steady by vaseline in saline solution, a similar single focal plane across all genotypes was recorded for 5 minutes to trace the puncta using Nikon C2 microscope with a 40X water objective. The puncta trafficking percentage and speed were analyzed by Imaris software (Bitplane, Zurich, Switzerland) with an Autoregressive Motion mode. The puncta-organelle contacting duration were calculated manually.

### Fly locomotion analysis

The rapid iterative negative geotaxis (RING) assay was used to analyze fly climbing ability as described previously [33, 34]. In brief, flies of different ages were collected and placed in vials with fly food (no yeast) for 1 day before being transferred to the cylinders for experiments. A total of 100 flies/genotype were analyzed in ten cylinders (inner diameter: 20 mm; height: 140 mm) placed side by side, and each contains ten unisex flies. Six initial mechanical impacts were applied so that all flies were slammed to the bottom of the cylinder and synchronized. Climbing distances for each fly were measured and averaged by RflyDetection software, which allows automatic detection of fly position within the cylinder using video images captured every 5 s (Sony digital camera, HDR-CX220E). At least three independent experiments were performed. Data were shown as mean ± SEM.

### Statistical analysis

Original images were imported into ImageJ (National Institutes of Health) for puncta number analyses. The intensity threshold of the relevant channel was set so that the maximum number of dots were selected without miscounting two adjacent dots as one. Original images were imported and automatically analyzed for colocalization using the colocalization plugin with the intensity correlation tool, resulting in Pearson’s R value (no threshold). GraphPad Prism 8 was used to display data in a scatter plot with bar form and calculate statistical significance. The Shapiro-Wilk test was used to check data distribution; for normally distributed datasets, the two-tailed t-test or ordinary one-way ANOVA followed by multiple comparisons test was used; and for datasets that did not have a normal distribution, the Mann Whitney test or Kruskal-Wallis tests were used.

## Results

### *atg9* is highly expressed in adult fly brain glia

To determine if Atg9 plays a role in glial cells, we first labeled adult fly brain glia by expressing *UAS-mCD8*.*GFP* under the control of a pan-glial driver *repo-GAL4* (*repo>mCD8*.*GFP*). These labeled glial cells were sorted by fluorescence-activated cell sorting (FACS) and analyzed for *atg9* expression by qRT-PCR (Figures 1A and 1B). These sorted GFP-positive cells exhibited high *repo* (a glial-specific marker) and low *elav* (a neuron-specific marker) expression, indicating that they are purified glial cells. Interestingly, *atg9* expression was considerably higher in GFP-positive cells than in GFP-negative cells (Figures 1B), suggesting that Atg9 might play a role in glia.

**Figure 1.**
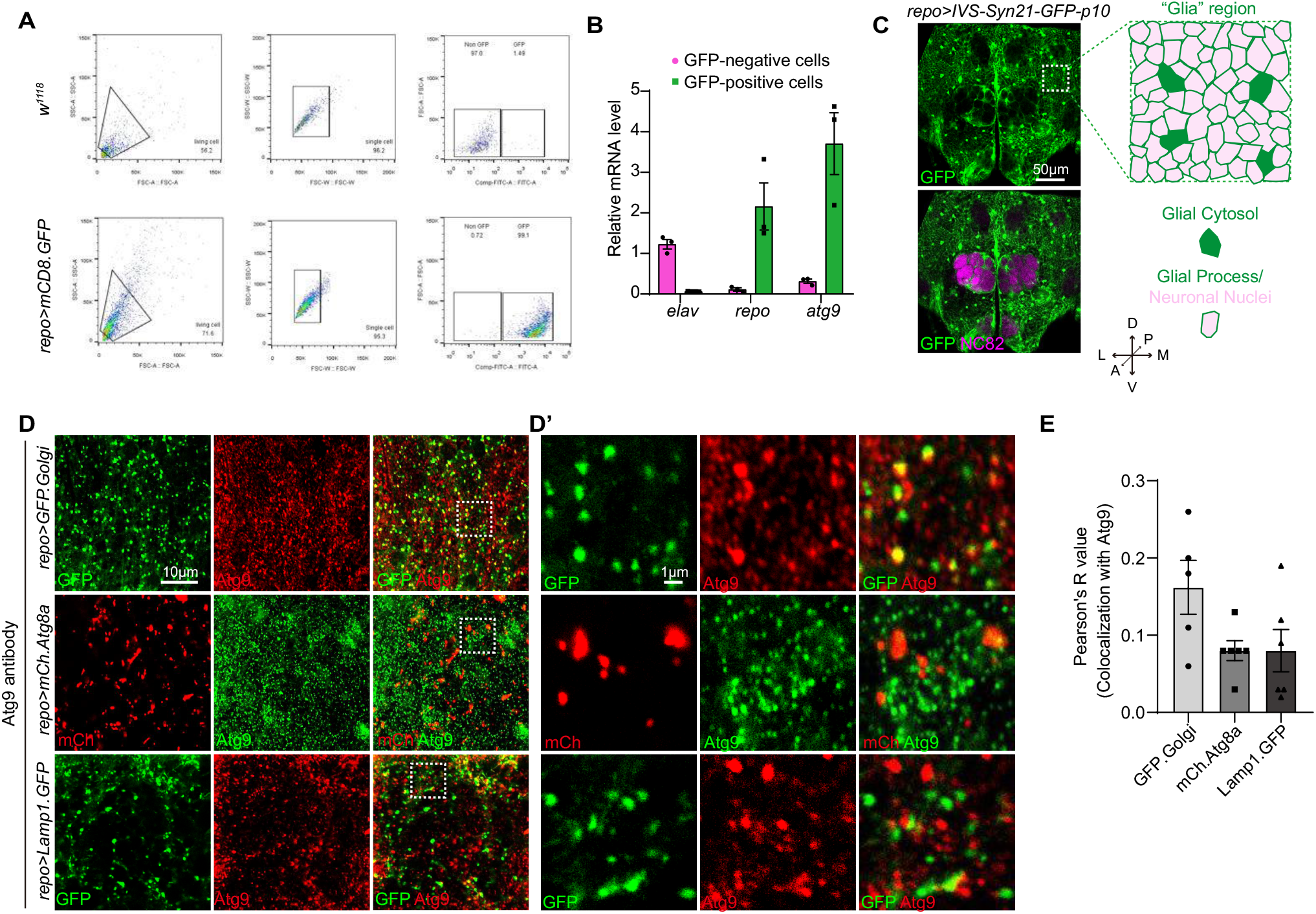
The expression and subcellular localization of Atg9 in adult fly brain glia. **(A and B)** GFP-positive glia were sorted by FACS analysis from flies expressing *UAS-mCD8*.*GFP* under the control of *repo-GAL4* (*repo>mCD8*.*GFP*). Cell sorting diagram (A) showed a distinct cell population representing GFP-positive glia. *atg9* expression level in sorted glia was analyzed by qRT-PCR (B). **(C)** Representative images of adult fly brains with glial membrane and nuclei labeled by *repo-GAL4* driven *UAS-IVS-Syn21-GFP-p10* (green), and axon neuropils labeled by the anti-NC82 antibodies (magenta). The orientation of the adult fly brain is depicted in a coordinate system (A: anterior, P: posterior, D: dorsal, V: ventral, M: medial, L: lateral). Dashed square in white indicates a “Glia” region at the dorsal-anterior part of the brain assessed throughout the study, and this region contains glial cytosol (solid green) and glial process (green line) wrapping neuronal nuclei (solid pink). **(D and E)** The subcellular location of endogenous Atg9 in glia was detected by anti-Atg9 antibodies. *repo>GFP*.*Golgi, repo>mCh*.*Atg8a*, and *repo>Lamp1*.*GFP* were used to label glial TGN, autophagosome, and lysosome, respectively. White dashed square in (D) and enlarged in (D’). (E) Quantification of Atg9 colocalization with different organelles as shown by the Pearson’s R value. Note that the value is higher for Atg9-TGN colocalization. Scale bars are indicated in the Figures. A serial confocal Z-stack sections were taken with 0.4 μm each, and the representative single layer images acquired at the similar plane of brains across all genotypes are shown. The data are present as mean SEM. Detailed fly genotypes are listed in Supplementary Materials.

### Atg9 is localized to TGN, autophagosome, and lysosome in adult fly brain glia

Next, the adult *Drosophila* brain was used as a system and analyzed by immunostaining analysis with different markers. The overall structure and location of different brain cell types were characterized with the anti-NC82 antibodies labeling the neuropile regions (magenta) and the glia-specific *repo-GAL4* driven UAS-*IVS-Syn21-GFP-p10* (*repo>IVS-Syn21-GFP-p10*) labeling the glial membrane and nuclei (green, Figure 1C). Our imaging analysis showed that glial cells are widely distributed, and the terminal processes of glia are often in close proximity to axon neuropils in the adult fly brains. A “Glia” region at the dorsal-anterior part of the brain, which contains both glial and neuronal components, was chosen for further analysis throughout the study.

During *Drosophila* development, Atg9 co-localizes with Atg8a autophagosomes, Rab11 endosomes, and Lamp1 endosomes-lysosomes in fat body [35]. We then checked whether Atg9 has a similar subcellular location in the adult fly brain glia. Transgenic flies such as *UAS-GFP*.*Golgi, UAS-mCherry*.*Atg8a*, and *UAS-Lamp1*.*GFP* were expressed by *repo-GAL4* and used to mark TGN, autophagosomes, and lysosomes in glia, respectively, while endogenous Atg9 was detected with the anti-Atg9 antibodies which specificity was validated by the *atg9*-RNAi (Figures S1A and S1B). Consistent with previous studies revealing Atg9 localization in other tissues, Atg9 also localizes on TGN, autophagosomes, and lysosomes in adult fly glia, with a higher portion of the protein localizing to TGN (Figures 1D and 1E).

Alternatively, a transgenic fly line expressing EGFP.Atg9 (*UAS-EGFP*.*Atg9*) was created for detecting Atg9 expression. EGFP.Atg9 expression and its colocalization with TGN, autophagosomes, or lysosomes were analyzed using different markers labeling these organelles. Under an overexpression condition, EGFP.Atg9, both in the glial cytosol and on the glial processes, showed significant colocalization with TGN, autophagosomes, and lysosomes (Figures 2A and 2A’). Taken together, these results indicate that glial Atg9 might play a role in autophagy as in other tissues.

**Figure 2.**
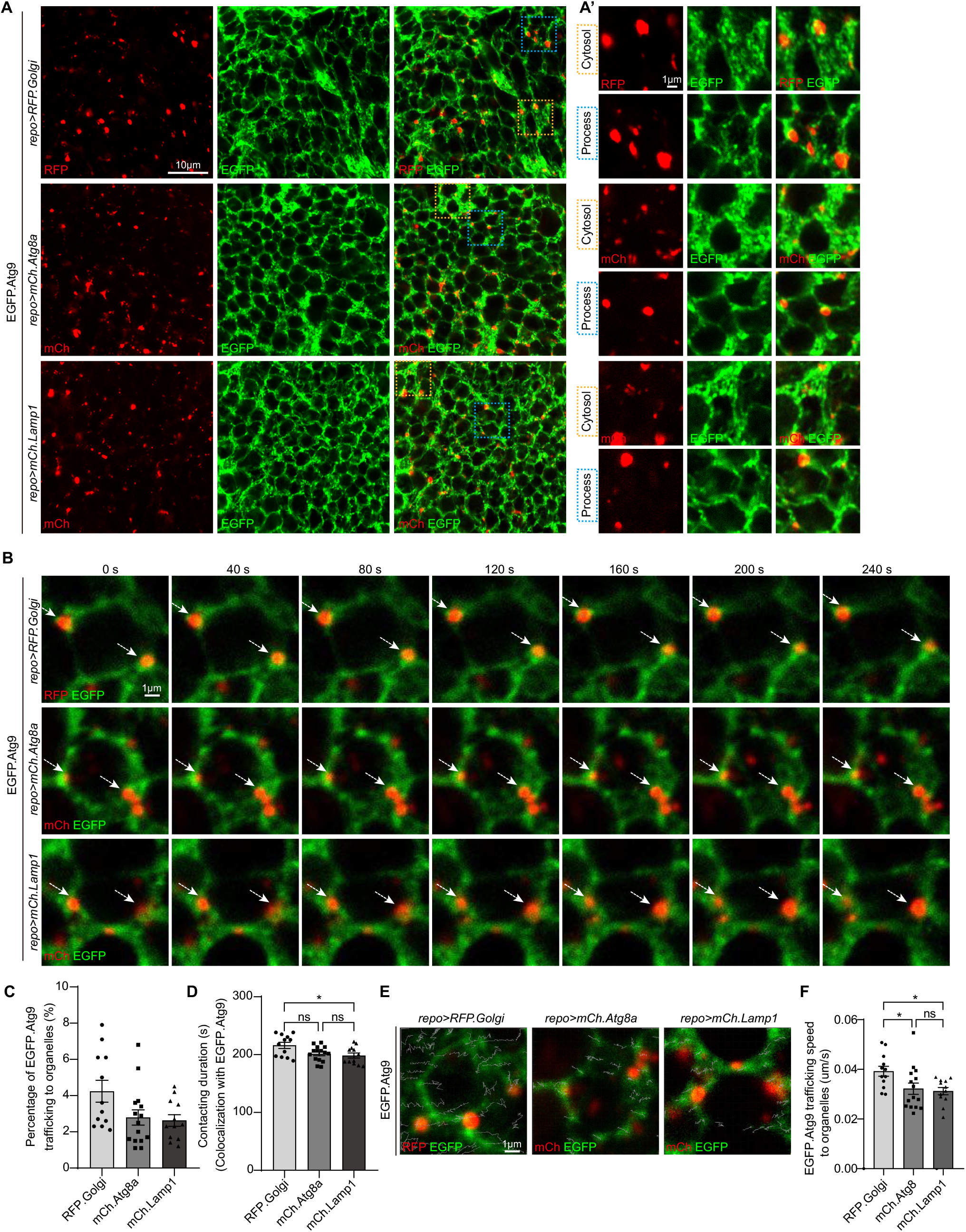
Trafficking dynamics of Atg9 in glial cells. **(A)** The subcellular location of Atg9 was detected with a transgenic fly expressing *UAS-EGFP*.*Atg9*. EGFP.Atg9 signals in both glial cytosol (yellow dashed square) and glial process (blue dashed square) were shown. *repo>RFP*.*Golgi, repo>mCh*.*Atg8a*, and *repo>mCh*.*Lamp1* were used to label glial TGN, autophagosome, and lysosome, respectively. A serial confocal Z-stack sections were taken with 0.4 μm each, and representative single layer images acquired at the similar plane of brains across all genotypes are shown. **(B-F)** Live-cell imaging (B and E) and quantifications (C, D and F) of EGFP.Atg9 trafficking dynamics in adult fly brain glia. A similar single focal plane across all genotypes was recorded and representative EGFP.Atg9 signals in glial process were shown. The percentage of EGFP.Atg9 puncta trafficking to the vicinity of different organelles (C), the duration of EGFP.Atg9 puncta contacting with different organelles (D), and the speed of EGFP.Atg9 puncta trafficking to different organelles (F) were analyzed. Scale bars are indicated in the Figures. The data are presented as mean SEM. P-values are determined by ordinary one-way ANOVA followed by Tukey’s multiple comparisons test (indicated with asterisks, ns no significance, ^*^p<0.05, ^**^ p<0.01, and ^***^ p<0.001).

### Glial Atg9 is trafficked to glial TGN, autophagosome, and lysosome

To further analyze the Atg9 dynamics on different organelles, live-cell imaging analysis of adult fly glia expressing EGFP.Atg9 in the presence of marker-labeled TGN, autophagosomes, or lysosomes was conducted (Figure 2B and Supplementary Videos 1-3). The percentage of EGFP.Atg9 trafficking to the vicinity of these organelles, their contacting time, and the speed were quantified. Notably, the percentage of EGFP.Atg9 puncta being trafficked to the vicinity of Golgi-positive puncta was higher, and the duration of Atg9-Golgi contact was also longer than the numbers for Atg9 trafficking to autophagosomes or lysosomes (Figures 2B-2D). Meanwhile, based on the trafficking route of EGFP.Atg9 puncta (Figure 2E), the speed of these puncta trafficking to TGN, autophagosomes, or lysosomes was calculated separately. Interestingly, Atg9 is most mobile when trafficking to TGN (Figures 2E and 2F). These observations revealed that Atg9 is trafficked to TGN, autophagosomes, or lysosomes in glia, and more Atg9 is trafficked to TGN in a faster speed.

### Lack of glial *atg9* decreases the number of omegasome and autophagosome

It has been shown that ATG9/Atg9 regulates early steps of autophagy, and is the main membrane source for the formation of omegasome and autophagosome [24, 27, 35]. To determine if Atg9 functions similarly in glia, GFP.ZFYVE1-labeled omegasomes [32] and mCh.Atg8a-labeled autophagosomes were quantified when inhibiting *atg9* expression in glia by RNAi (either *atg9*-RNAi BL28055 or *atg9*-RNAi^#2^ V10045). Flies expressing *UAS-LacZ* was used as a control. The RNAi knockdown efficiencies were validated by qRT-PCR, Western blot (WB), and immunostaining (Figures 3A-3C, S1A-S1C). Notably, lack of glial *atg9* caused a significant reduction on the number of omegasome (Figures 3D and 3E) and autophagosome (Figures 3F and 3G, Figures S1D and S1E). These results indicate that glial Atg9 functions in autophagy by affecting the number of early autophagic structures such as omegasome and autophagosome.

**Figure 3.**
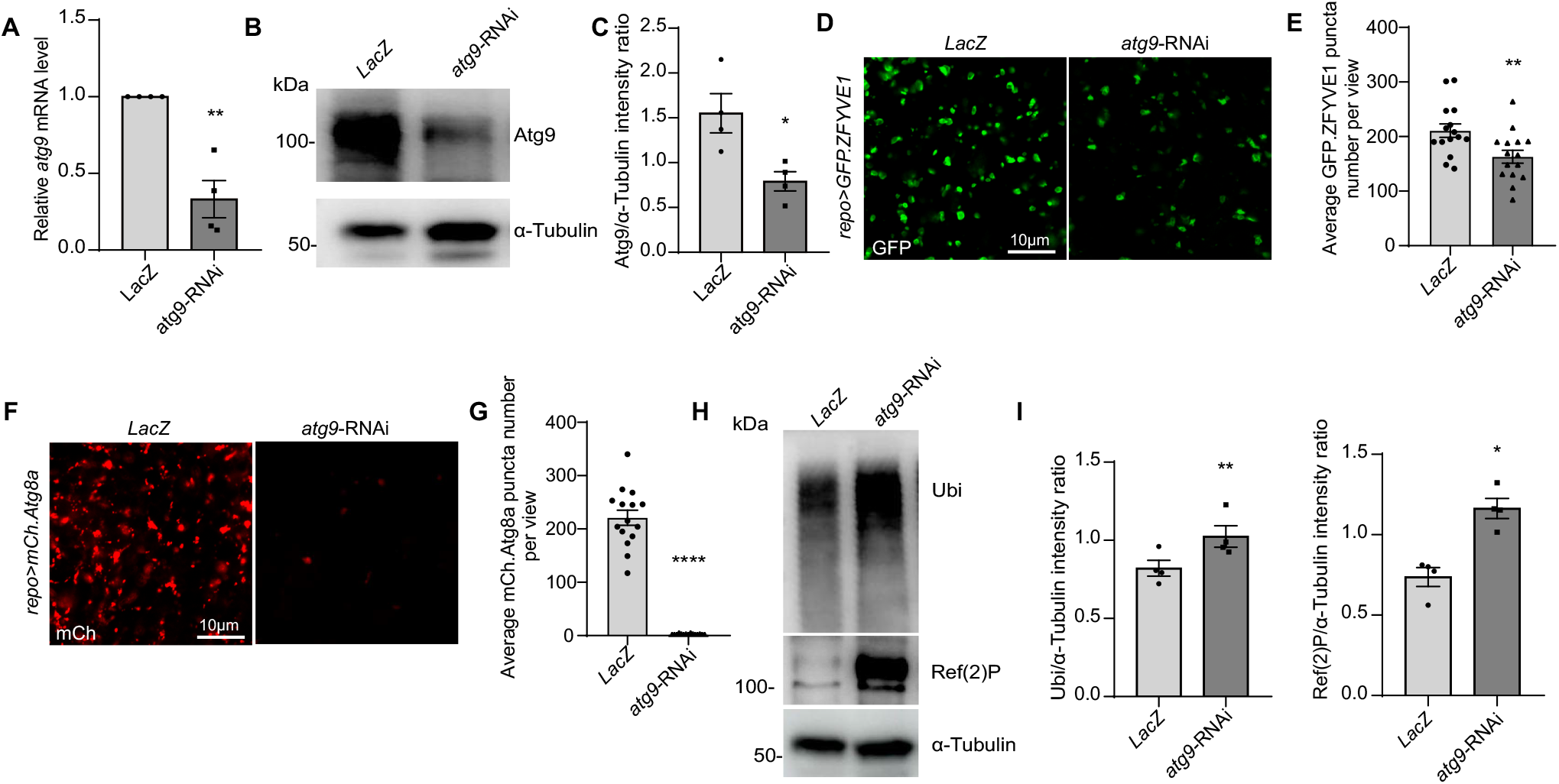
Glial Atg9 regulates omegasome and autophagosome formation and substrates degradation. **(A-C)** The knocking down efficiency of *atg9*-RNAi lines were analyzed by qRT-PCR (A) and Western blot (B and C). Adult fly heads from the control and *repo>atg9-RNAi* strains were collected for RNA and protein samples. *rp49* and α-Tubulin serve as controls for qRT-PCR and WB, respectively. Note that both mRNA and protein levels of Atg9 decreased when expressing *atg9*-RNAi in glia. **(D-G)** Representative images (D and F) and quantifications (E and G) of GFP.ZFYVE1- and mCh.Atg8a-positive puncta in control and *repo>atg9-RNAi* adult fly brains. Note that depletion of glial *atg9* reduces both omegasome and autophagosome number. **(H and I)** Autophagic substrates degradation, such as Ref(2)P (the *Drosophila* P62 homolog) and Ubiquitin (Ubi), were impaired in *repo>atg9-RNAi* adult fly brains. Scale bars are indicated in the Figures. A serial confocal Z-stack sections were taken with 0.4 μm each, and representative single layer images acquired at the similar plane of brains across all genotypes are shown. The data are presented as mean SEM. P-values are determined using the two-tailed unpaired t-test or Mann-Whitney test (indicated with asterisks, ns no significance, ^*^ p<0.05, ^**^ p<0.01, and ^***^p<0.001).

### Lack of glial *atg9* blocks autophagic substrate degradation

Given that Atg9 regulates the number of omegasome and autophagosome, we next sought to investigate if the Atg9-mediated mechanism affects the efficiency of glial autophagy. Results from both WB and immunostaining analysis indicated that the levels of autophagic substrates such as Ref(2)P, the *Drosophila* ortholog of P62, and ubiquitin (Ubi) accumulated significantly when depleting glial *atg9* (Figures 3H and 3I, S1F and S1G), suggesting that Atg9 contributes to autophagic substrate degradation in glia.

### Lack of glial *atg9* leads to parkinsonian symptoms

As ATG9 dysregulation has been implicated in PD [27, 28], we then assessed whether these aforementioned Atg9-mediated autophagic defects in glia plays a role in PD using a *Drosophila* PD model [34]. DA neurodegeneration was detected by the anti-Dopa decarboxylase (DDC) antibodies (Figure 4A). Notably, lack of glial *atg9* caused age-dependent DA neuron loss at the PPM1/2 cluster, which have been shown to degenerate upon α-syn overexpression [36]. The DA neuron number was not affected by the absence of glial *atg9* immediately after eclosion, ruling out the developmental defects (Figures 4B and 4C). Furthermore, fly climbing ability were quantified by an automated rapid iterative negative geotaxis (RING) assay [33, 34], and lack of glial *atg9* caused significant locomotion defects in an age-dependent manner (Figures 4D). Taken together, our results demonstrate that glial Atg9 contributes to DA neurodegeneration and locomotor function, both typical hallmarks of PD [34].

**Figure 4.**
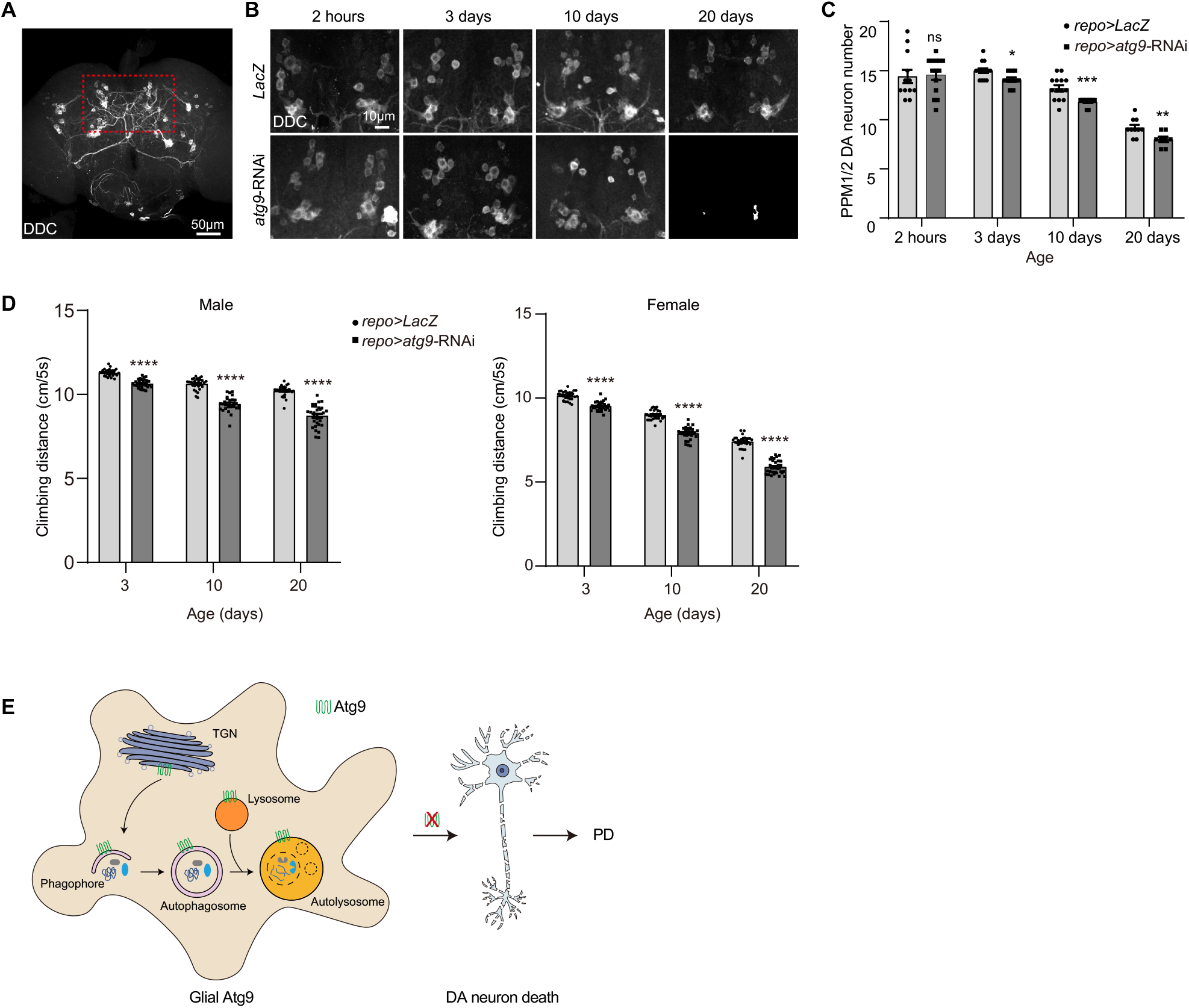
Glial Atg9 contributes to parkinsonian symptoms. **(A)** Illustration of an adult fly brain with different DA neuron clusters. The selected PPM1/2 clusters are enclosed by the red dotted square and enlarged in B. **(B and C)** Representative images (B) and quantification (C) of PPM1/2 DA neuron number in adult fly brains with the indicated genotypes. Note that the number of DA neurons at the PPM1/2 cluster decreases significantly with age in the absence of glial *atg9*. **(D)** Lacking glial *atg9* caused age-dependent locomotion defects in both male and female flies. **(E)** A model illustrating the subcellular location of glial Atg9, and the link between the glial function of Atg9 and PD. Scale bars are indicated in the Figures. A serial confocal Z-stack sections were taken with 0.4 μm each, and images are shown as maximal projection. The data are presented as mean SEM. P-values are determined using the two-tailed unpaired t-test, Mann-Whitney test (indicated with asterisks, ns no significance, ^*^ p<0.05, ^**^p<0.01, and ^***^ p<0.001).

## Discussion

In the present study, we investigated the glial function of the autophagy protein Atg9 in the context of PD (Figures 4E). *atg9* is highly expressed in adult fly glia and diversely localized in TGN, autophagosomes, and lysosomes in glia. In addition, Atg9 is persistently in contact with these organelles. Consistent to Atg9 function in fat body, Atg9 functions in glia by regulating the formation of omegasome and autophagosome, hence the degradation of autophagic substrates. Importantly, loss of glial *atg9* induces parkinsonian symptoms including progressive DA neuron loss and locomotion deficits. Thus, our findings suggest that Atg9 modulates autophagic function in glia and contributes to PD.

In the past decades, numerous studies have demonstrated that glial activation and neuroinflammation are key features for PD onset and progression [6, 8, 9]. Neuronally released α-syn activates astrocytes or microglia, leading to either a neuroprotective or neurotoxic feedback from glia to neurons [9, 37]. Many PD risk genes are also highly expressed in glial cells [7, 38-40]. These evidence demonstrate the importance of glia in PD. Interestingly, our results indicate that the glial function of Atg9 contributes to parkinsonian symptoms in a *Drosophila* PD model that we previously established [34]. Consistent to previous results revealing that *atg9* null mutant flies exhibit shortened lifespan, locomotor defects, and increased susceptibility to stresses [41], our findings show that lacking glial *atg9* induces age-dependent progressive DA neuron loss and locomotion defects. In addition, knocking down *atg9* in glia leads to impaired formation of autophagic precursors and substrate degradation, potentially leading to protein aggregate accumulation, a key pathological hallmark of neurodegenerative diseases. Thus, we confirm the autophagic function of Atg9 in glia and link the function to PD. These findings underscore the importance of glial regulatory mechanism in PD and shed lights on the potential interest of a glial autophagy protein as a therapeutic target.

## Acknowledgments

We thank Bloomington *Drosophila* Stock Center, Vienna *Drosophila* RNAi Center, the Core Facility of *Drosophila* Resource and Technology, Shanghai Institute of Biochemistry and Cell Biology, Chinese Academy of Sciences, Developmental Studies Hybridoma Bank, Chao Tong, and Gerald M Rubin for fly stocks and antibodies. We also thank the Molecular Imaging Core Facility (MICF) and the Molecular and Cell Biology Core Facility (MCBCF) at the School of Life Science and Technology, ShanghaiTech University for providing technical support; Ho lab members for discussion and comments. This work was supported by grants from ShanghaiTech and National Natural Science Foundation of China (31871039 and 32170962).

## Author contributions

S.Y, M.S.H, and S.Z conceived and designed the study. S.Y, L.W, and S.Z performed the experiments. S.Y, M.S.H, and S.Z analyzed the data and wrote the paper. All authors read and approved the manuscript.

