## Supplementary figures and images for "The autophagy protein Atg9 functions in glia and contributes to parkinsonian symptoms in a *Drosophila* model of Parkinson’s disease"

### Supplementary Figure 1

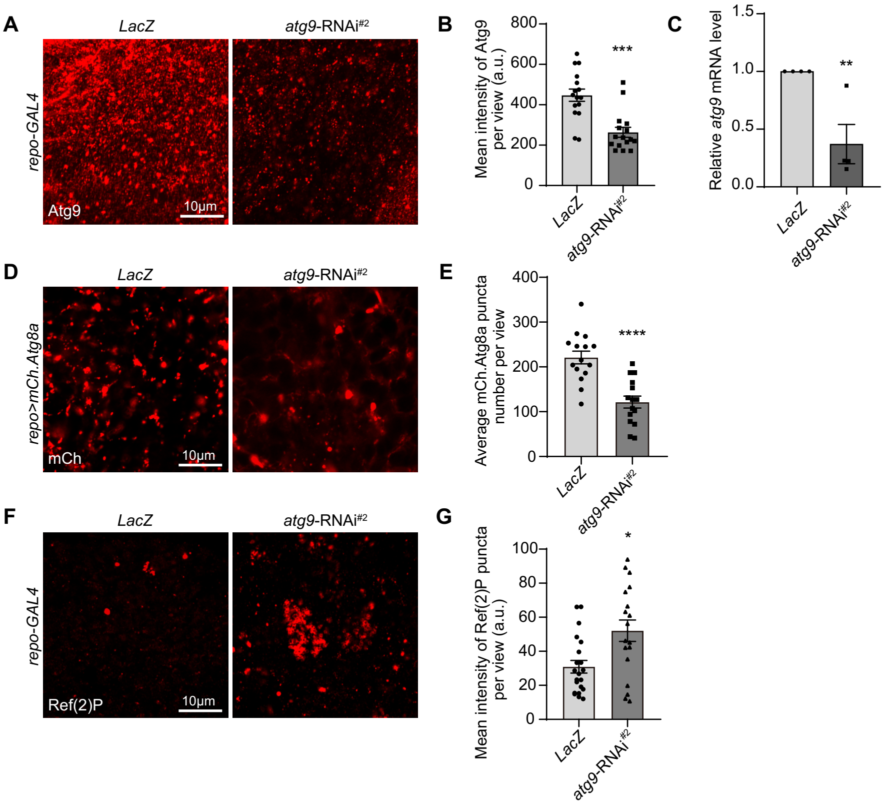
