## Supplementary Materials for "The autophagy protein Atg9 functions in glia and contributes to parkinsonian symptoms in a *Drosophila* model of Parkinson’s disease"

**This PDF file includes:**

Supplementary Figure S1

Supplementary Video Legend

*Drosophila* Genotype Table

**Supplementary Figure**

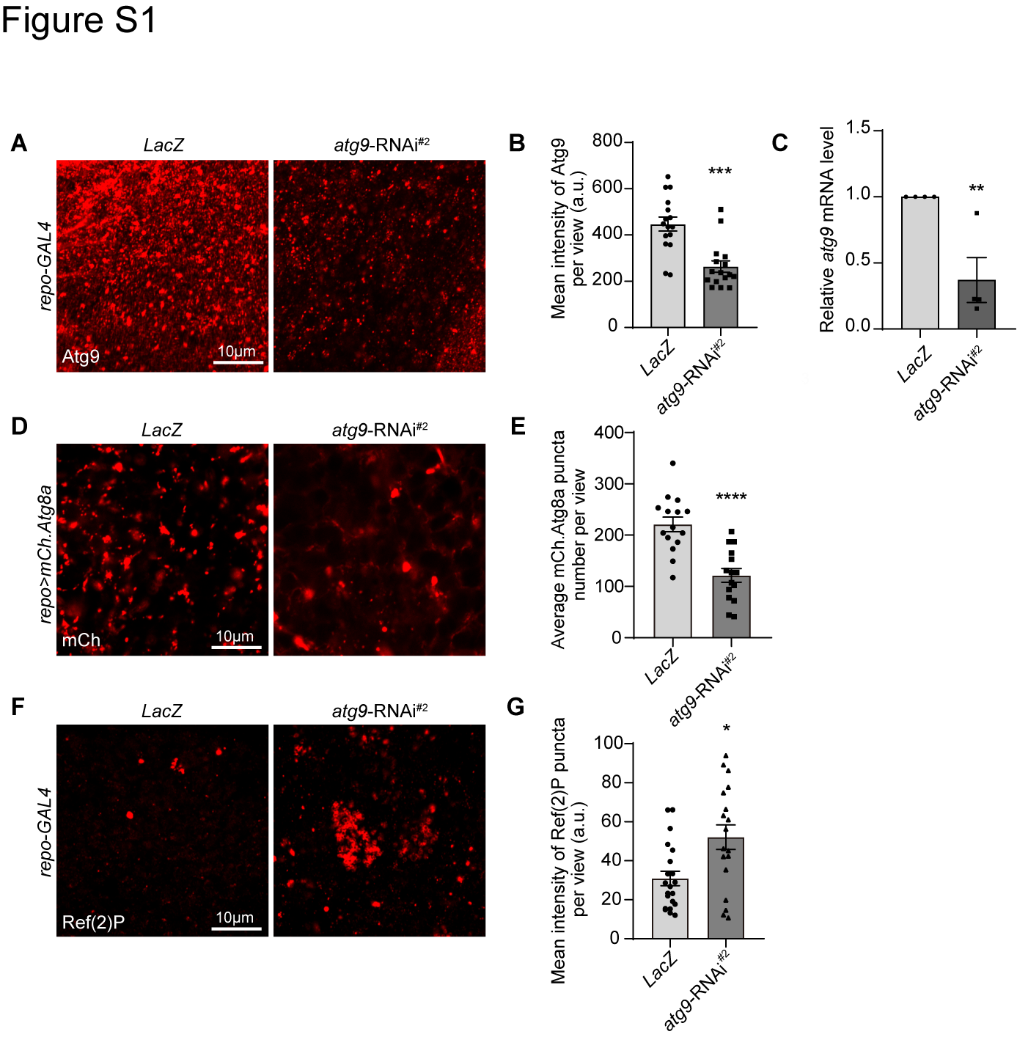

**Figure S1 (related to Figure 3). Expressing *atg9-*RNAi^#2^ in glia impairs autophagosome formation and substrates degradation.**

**(A-C)** The knocking down efficiency of *atg9*-RNAi lines was analyzed by immunostaining with anti-Atg9 antibodies (A and B) and qRT-PCR (C). **(D-G)** Representative images (D and F) and quantifications (E and G) of glial autophagosome number and anti-Ref(2)P staining in the control and *repo>atg9-RNAi^#2^* adult fly brains. Note that similar to the other RNAi, expressing *atg9*-RNAi^#2^ in glia also decreased autophagosome number and impairs autophagic substrate degradation. Scale bars are indicated in the Figures. The data are presented as mean SEM. P-values are determined using the two-tailed unpaired t-test or Mann-Whitney test, (indicated with asterisks, ns no significance, * p<0.05, ** p<0.01, and *** p<0.001).

**Supplementary Video Legend**

**Videos. Live-cell imaging of Atg9 trafficking**

Live-cell imaging of EGFP.Atg9 trafficking to RFP.Golgi (Video 1), mCh.Atg8a (Video 2), and mCh.Lamp1 (Video 3) in the “Glia” region. Adult fly brains were dissected and kept in saline solution, and a similar single focal plane across all genotypes was recorded.

***Drosophila* Genotype Table**

| **Fig. 1** |  |
| --- | --- |
| (A and B) | *w^1118^* |
|  | *repo-GAL4, UAS-mCD8.GFP/+* |
| (C) | *repo-GAL4, 10xUAS-IVS-Syn21-GFP-p10/+* |
| (D and E) | *UASp-GFP.Golgi/+; repo-GAL4/+* |
|  | *UASp-mCherry.Atg8a/+; repo-GAL4/+* |
|  | *UAS-Lamp1.GFP/+; repo-GAL4/+* |
| **Fig. 2** |  |
| (A-F) | *UASp-RFP.Golgi/+; repo-GAL4, UAS-EGFP.Atg9/+* |
|  | *UASp-mCherry.Atg8a/+; repo-GAL4, UAS-EGFP.Atg9/+* |
|  | *UAS-mCherry.Lamp1/+; repo-GAL4, UAS-EGFP.Atg9/+* |
| **Fig. 3** |  |
| (A-C) | *repo-GAL4/UAS-LacZ* |
|  | *repo-GAL4/UAS-atg9-*RNAi |
| (D and E) | *UAS-GFP.ZFYVE1/+; repo-GAL4 UAS-mCD8.RFP/UAS-LacZ* |
|  | *UAS-GFP.ZFYVE1/+; repo-GAL4 UAS-mCD8.RFP/UAS-atg9-*RNAi |
| (F and G) | *UASp-mCherry.Atg8a/+; repo-GAL4/UAS-LacZ* |
|  | *UASp-mCherry.Atg8a/+; repo-GAL4/UAS-atg9-*RNAi |
| (H and I) | *repo-GAL4/UAS-LacZ* |
|  | *repo-GAL4/UAS-atg9-*RNAi |
| **Fig. 4** |  |
| (A) | *w^1118^* |
| (B-D) | *repo-GAL4/UAS-LacZ* |
|  | *repo-GAL4/UAS-atg9-*RNAi |
| **Fig. S1** |  |
| (A-C) | *repo-GAL4/UAS-LacZ* |
|  | *repo-GAL4/UAS-atg9-*RNAi^#2^ |
| (D and E) | *UASp-mCherry.Atg8a/+; repo-GAL4/UAS-LacZ* |
|  | *UASp-mCherry.Atg8a/+; repo-GAL4/UAS-atg9-*RNAi^#2^ |
| (F and G) | *repo-GAL4/UAS-LacZ* |
|  | *repo-GAL4/UAS-atg9-*RNAi^#2^ |
